# Effects of Human-Animal Interaction on Dog Salivary and Plasma Oxytocin and Vasopressin

**DOI:** 10.1101/151506

**Authors:** Evan L. MacLean, Laurence R. Gesquiere, Nancy Gee, Kerinne Levy, W. Lance Martin, C. Sue Carter

## Abstract

Oxytocin (OT) and Vasopressin (AVP) are neuropeptides with diverse effects on social behavior, cognition and stress responses. Recent studies suggest that OT facilitates and responds to affiliative forms of human-animal interaction (HAI). However, previous studies measuring OT and AVP in dogs have been limited to measures from blood or urine, which present concerns related to the invasiveness of sample collection, the potential for matrix interference in immunoassays, and whether samples can be collected at precise time points to assess event-linked endocrine responses. Previous studies from our laboratory validated salivary measures of OT and AVP in dogs, however, it is currently unknown whether these measures respond dynamically to aspects of HAI. Here, we investigated the effects of affiliative forms of HAI on both plasma and salivary OT and AVP in dogs. We employed a between-subjects design with a group of Labrador retrievers and Labrador retriever X golden retriever crosses (23 females, 15 males). Half of the dogs engaged in 10 minutes of free-form friendly interaction with a human experimenter (HAI condition), and the other half rested quietly in the same environment, without human interaction (control condition). We collected blood and saliva samples before, and immediately following both experimental conditions, and all samples were analyzed using enzyme-linked immunosorbent assays (ELISAs) following previously validated protocols. Dogs participating in HAI exhibited a significant increase in both salivary OT (+39%) and plasma OT (+5.7%) whereas dogs in the control group did not. Salivary AVP showed no change in the HAI group but increased significantly (+33%) in the control group. Plasma AVP decreased significantly following HAI (-13%) but did not change across time in the control condition. Within the dogs exposed to HAI, increases in salivary OT, and decreases in plasma AVP, were predicted by the extent of affiliative behaviour between the dog and human (indexed by scores from a principal components analysis of social behaviours between the dog and human). Collectively our results suggest that measures of salivary OT and AVP provide useful biomarkers in studies of HAI, and afford a flexible and noninvasive toolkit than can be employed in diverse research contexts.

---

Studies throughout the last three decades have explored the psychological and physiological effects of human-animal interaction (HAI). Often the aim of such studies is to characterize the mechanisms through which nonhuman animals affect human health and wellbeing, and in turn, how interaction with humans affects animal participants. Recently such studies have begun to focus on the neuropeptide oxytocin (OT), which, together with (structurally related) arginine vasopressin, is well known for its roles in facilitating selective social bonds, and regulating various aspects of social behavior and cognition in mammals (Carter, Grippo, Pournajafi-Nazarloo, Ruscio, & Porges, 2008). For example, central OT administration can facilitate maternal behavior in sexually naïve rodents (Caldwell & Young III, 2006; Pedersen & Prange, 1979), and both OT and AVP are critical to the formation of partner preference in pair-bonded species (Carter, Williams, Witt, & Insel, 1992). With regard to the establishment of selective social attachment, the OT receptor (OXTR) is highly expressed in the nucleus accumbens of some pair-bonded species, suggesting that OT may help to encode social reward through modulation of the mesolimbic dopamine pathway (Insel & Shapiro, 1992; Lim & Young, 2006).

Research on the role of OT in HAI has been conducted by measuring OT release during interactions between humans and dogs, and evaluating the effects of exogenous OT administration on dog behavior in this context. Studies of endogenous OT have revealed increases in blood or urinary OT concentrations in both humans and dogs following affiliative interaction in dog-human dyads (Handlin et al., 2011; Miller et al., 2009; Nagasawa, Kikusui, Onaka, & Ohta, 2009; Nagasawa et al., 2015; Rehn, Handlin, Uvnäs-Moberg, & Keeling, 2014). Additionally, preliminary evidence suggests that dogs bred for friendly and non-aggressive temperaments are characterized by high levels of plasma OT, relative to pet dogs (MacLean, Gesquiere, Gruen, et al., Submitted). Studies using intranasal OT administration in dogs suggest that exogenous OT can facilitate social play (Romero, Nagasawa, Mogi, Hasegawa, & Kikusui, 2015), and promote bonding with both conspecifics and humans (Romero, Nagasawa, Mogi, Hasegawa, & Kikusui, 2014). Additionally, recent studies have demonstrated that exogenous OT can improve dogs’ sensitivity to human communication (Macchitella et al., 2016; Oliva, Rault, Appleton, & Lill, 2015). Thus, a rapidly growing body of research suggests that OT pathways may be centrally involved in affiliative forms of HAI (Beetz, Uvnäs-Moberg, Julius, & Kotrschal, 2012; Carter & Porges, 2016; MacLean & Hare, 2015).

We are not aware of any studies investigating AVP in the context of HAI, however, many of the effects of OT are also critically dependent on AVP (Carter, 1998; Landgraf & Neumann, 2004). Although AVP plays important roles in selective sociality, particularly among males (Caldwell, Lee, Macbeth, & Young III, 2008), its effects are often antagonistic to those of OT. For example, many of the social effects of OT during HAI may be facilitated by OT’s attenuation of sympathetic arousal (Buttner, 2016; Kis, Kanizsár, Gácsi, & Topál, 2014) through actions in the hypothalamus (Dabrowska et al., 2011), and on the vagus nerve (Porges, 2003; Porges, 2007; Porges, 2011). In contrast, AVP activates the hypothalamic-pituitary-adrenal (HPA) axis, and is more strongly linked to anxiety and aggression (Coccaro, Kavoussi, Hauger, Cooper, & Ferris, 1998; Neumann & Landgraf, 2012). We are aware of two studies investigating relationships between AVP and behavior in dogs, both of which revealed positive associations with anxiety or aggression (Hydbring-Sandberg et al., 2004; MacLean, Gesquiere, Gruen, et al., Submitted). Given its structural and functional relationships with OT, we expect that AVP may also regulate or respond to aspects of HAI, although these effects are likely to differ from those for OT.

To date, all studies of OT and AVP in dogs engaged in HAI have quantified peptide concentrations in blood or urine samples. Although urine sampling can be performed non-invasively, it yields poor temporal resolution and characterizes long periods of peptidergic activity. In contrast, blood sampling can capture acute changes in peptide release, but is an invasive procedure that may induce stress and acute pain, and is not well suited to many of the contexts in which HAI studies are conducted. Additionally, OT and AVP rapidly bind to other molecules in blood (and likely urine) which can lead to matrix interference in assays, or erroneously low estimates of peptide concentrations (Brandtzaeg et al., 2016; Martin & Carter, 2013). We recently validated methods for quantifying OT and AVP in dog saliva samples, which can be collected noninvasively, and at precise time points during HAI. Enzyme-linked immunosorbent assays (ELISAs) yielded good parallelism and accuracy, and did not require an extraction procedure. Lastly, we previously measured salivary OT in nursing dams and detected an acute rise in OT associated with milk letdown, providing an initial biological validation of this measure (MacLean, Gesquiere, Gee, et al., Submitted).

In the current studies, we evaluated both plasma and salivary OT and AVP concentrations in dogs, before and after affiliative interaction with a human, or in a control condition. Because the time course of salivary OT/AVP release during HAI was unknown, we first conducted a short pilot study to identify time points associated with changes in salivary OT/AVP concentrations during HAI. We then conducted an experiment with dogs assigned to an HAI or control condition, and assessed OT/AVP changes over time in both groups, and as a function of behavior during the test period. Based on the studies described above, we hypothesized that dogs in the HAI condition would exhibit increases in OT across the study period, and that any such changes would be larger in the HAI group than the control group. We also hypothesized that within the HAI group, changes in OT would be predicted by the extent of affiliative behavior between the human and the dog during the study. Because no studies have investigated effects of HAI on AVP, we had no specific hypotheses for the nature of this response. However, given that AVP also plays important roles in social emotions and behavior, we expected that dogs in the HAI group would exhibit changes in AVP that differed from those of dogs in the control group.

## Pilot Study

Prior to the main HAI study, we conducted a short pilot study with 10 dogs to identify the optimal sampling periods for detecting changes in salivary OT and AVP. Based on our studies with nursing dams (MacLean, Gesquiere, Gee, et al., Submitted), and recent studies measuring salivary OT in humans (de Jong et al., 2015), we expected changes in saliva to be rapid. Therefore, we assessed both plasma and salivary OT and AVP at 5 and 10 minutes following the start of HAI.

## Method

### Subjects

We tested dog subjects from the breeding colony at Canine Companions for Independence (CCI) in Santa Rosa, CA, USA. The pilot sample included 10 dogs (8 female, 2 male, 4 Labrador retrievers, 6 Labrador retriever X golden retriever crosses, mean age = 1.7 years (range = 1.6-1.8 years)). All dogs were pair-housed in indoor-outdoor kennels with *ad libitum* access to water and daily access to large outdoor play yards. Subjects were tested in a quiet room inside a familiar building on CCI’s campus. Client consent was obtained for participation of all dogs and all animal procedures were approved by the Duke University IACUC (protocol #A138- 11-06).

### Procedure

Prior to the test each dog was allowed to rest quietly in a crate (outside the test room) for 30 minutes. The dog was then taken to a nearby room and we collected a baseline blood and saliva sample. Saliva samples were collected using the Salimetrics Children’s swab as described in MacLean et al. (Submitted). Blood samples were collected from the cephalic vein into vacutainers (3 mL) containing ethylenediaminetetraacetic acid (EDTA). Blood and saliva samples were collected concurrently to minimize the time required for these procedures. Specifically, at the start of the collection period, one experimenter placed the swabs between the dog’s cheek and mandibular teeth and gently held the dog’s mouth closed while the second experimenter performed the blood collection. All samples were immediately placed on ice after collection.

Following this initial sample, dogs were allowed to rest in a crate inside the test room for 5 minutes of rest prior to the start of the behavioral interaction. After 5 minutes, dogs were released from the crate and allowed to interact freely with the experimenter. The experimenter attempted to engage the dog in friendly interaction, including gently petting the dog, and speaking to the dog in a friendly tone while making eye contact. However, the experimenter allowed the dog to lead these interactions, and dogs were always free to disengage and move away from the experimenter at any point during the interaction. If subjects attempted to engage the experimenter in play (e.g. performing a play bow, chasing, or nuzzling the experimenter), the experimenter engaged with the dog in these more active forms of interaction. After 5 minutes of HAI we collected a second blood and saliva sample from the dog, and immediately resumed HAI for another five minutes.

The final blood and saliva samples were collected 10 minutes after the start of HAI and dogs received a food reward at the conclusion of this period (following the final saliva collection). The second and third blood samples were collected from (1) the cephalic vein on the opposite forelimb from the initial draw, and (2) the jugular vein, in order to avoid repeated needle punctures in the same location. We did not use a catheter for repeated collections from the same site because pilot tests revealed that catheters did not reliably maintain access to the vein when dogs were allowed to move freely between collection periods, and the procedures required to position, wrap, and access the catheter repeatedly were deemed more likely to cause discomfort for the dog than single collections from different sites. All biological samples were immediately frozen at -20 °C.

### Hormone analysis

All samples were analyzed by enzyme-linked immunoassay (ELISA) following protocols previously validated in our laboratory (MacLean, Gesquiere, Gee, et al., Submitted; MacLean, Gesquiere, Gruen, et al., Submitted). OT samples were measured using a commercially available kit from Cayman Chemical (Item #500440) and AVP samples were measured using a commercially available kit from Enzo Life Sciences (ADI-900-017A). Saliva samples were not extracted based on the results of validation studies (MacLean, Gesquiere, Gee, et al., Submitted) but plasma samples were processed using solid phase extraction (to isolate free peptide concentrations) with the protocols described in MacLean et al. (Submitted).

### Statistical analysis

All statistical analyses were performed in the R programming environment for statistical computing (R Core Team, 2017). For each matrix (saliva, plasma) and hormone (OT, AVP), we used linear mixed models to predict the log transformed hormone concentration as a function of a fixed effect of time (baseline, +5 mins, +10 mins) and a random effect of subject ID. We used planned Dunnett contrasts to assess mean differences between hormone concentrations at baseline and + 5 minutes, and between baseline and +10 minutes.

## Results and Discussion

Table 1 shows the results of linear mixed models predicting changes in hormone concentrations as a function of time. We inspected the data for time points associated with the largest deviations from baseline values.

**Table 1.** Pilot study results. Baseline (T0) was used as the reference value and the T5-T0 and T10-T0 Dunnett contrasts show the estimates, and standard errors of estimates, for changes in hormone concentrations at each time point, relative to T0 (β = change in log pg/mL from T0).

|  | T5 – T0 |  |  | T10 – T0 |  |  |
| --- | --- | --- | --- | --- | --- | --- |
| | $\beta$ | SE | p | $\beta$ | SE | p |
| Salivary OT | 0.14 | 0.17 | 0.40 | 0.33 | 0.17 | 0.05 |
| Plasma OT | 0.03 | 0.04 | 0.48 | 0.01 | 0.04 | 0.83 |
| Salivary AVP | -0.19 | 0.15 | 0.22 | -0.05 | 0.15 | 0.73 |
| Plasma AVP | -0.05 | 0.12 | 0.67 | -0.07 | 0.12 | 0.56 |

Salivary OT increased slightly from baseline to +5 minutes, with a further increase at +10 minutes that was marginally different from baseline (p = 0.05; Table 1). Changes in plasma OT and both salivary and plasma AVP were minimal, with no clear deviations from baseline. Therefore, we identified the +10-minute measure as the most likely to show an HAI related effect in salivary OT, and collected samples at this time point in Experiment 1.

## Experiment 1

### Method

#### Subjects

We tested 38 dogs (11 Labrador retrievers, 27 Labrador retriever X golden retriever crosses, 23 female, 15 male, mean age = 1.8 years, range = 1.6 - 2.2 years). All subjects were assistance dogs in training at Canine Companions for Independence and were housed and cared for as described above. Half of the subjects were assigned to the HAI condition, and half were assigned to the control condition (see below). Groups were matched closely based on sex and breed (Table 2).

**Table 2.** Subject demographics by condition for Experiment 1. HAI = Human-Animal Interaction; LAB = Labrador Retriever; LGX = Labrador X golden retriever cross

| Condition | Breed | Female | Male |
| --- | --- | --- | --- |
| HAI | LAB | 5 | 1 |
|  | LGX | 6 | 7 |
| Control | LAB | 4 | 1 |
|  | LGX | 8 | 6 |

#### Procedure

The HAI procedure was identical to that in the pilot study with the exception that we collected blood and saliva samples at only one time point (+10 minutes) after the baseline measure, based on preliminary data suggesting the greatest changes in salivary OT at this time. Therefore, the experimenter and dog had 10 continuous minutes of interaction prior to the post-test sample collection. In the control condition, dogs were placed inside an exercise pen (~3m X 3m) in the test room. The experimenter remained in the room with the dog but did not interact, speak to, or make eye contact with the dog during the 10-minute test period. As in the HAI condition, after 10 minutes we collected a post-test blood and saliva sample. After the study, dogs in both conditions received a food reward.

#### Behavioral coding

To assess whether specific behaviors or forms of social interaction were related to individual differences in hormonal response, we coded the duration (s) of several behaviors from video. Locomotion and postural variables were coded for subjects in both the control and HAI conditions, and consisted of the following: (1) *locomotion*: time walking, running, or jumping, with the onset of the behavior marked by three consecutive steps (to disambiguate locomotion from minor bodily repositioning), and of the offset of the behavior marked by a lack of movement for ≥ 1 s; (2) *upright*: time standing, walking, running, or jumping; (3) *lying (prone)*: time lying with stomach against the floor; (4) *lying (supine)*: time lying on back or side with stomach exposed; (5) *sitting*: time with rump on ground and forelegs extended.

For subjects in the HAI condition we also coded the following behaviors relating to interaction with the experimenter: (6) *physical contact*: time during which any part of the experimenter’s body was in physical contact with any part of the dog’s body; (7) *licking*: time in which the dog’s tongue was in contact with the experimenter’s body; (8) *play*: time in which the dog engaged in play bows, chasing, or gentle mouthing with the experimenter, with the onset marked by the first occurrence of any of these behaviors, and the offset marked by a period of ≥ 2 s without the dog engaging in any of these behaviors. A second independent rater scored all behaviors for random sample of ~50% of observations, and inter-rater reliability was excellent for all measures (Pearson correlation, mean: R = 0.997 (min = 0.993, max = 1.0)

#### Hormonal analysis

Plasma and salivary OT and AVP were assayed using the same ELISA kits and protocols employed in the pilot study. Inter-assay coefficients of variation were 11.1% (salivary OT), 4.2% (salivary AVP), 11.7% (plasma OT) and 11.2% (plasma AVP).

#### Statistical analysis

All statistical analyses were performed in the R programming environment for statistical computing (R Core Team, 2017). To compare changes in OT/AVP across time between conditions, we used linear mixed models with the log transformed OT/AVP concentrations predicted as a function of fixed effects for sex (male, female), time (pre, post), condition (control, HAI), the time X condition interaction, and a random effect for subject ID. This model was fitted separately with each endocrine measure as the dependent variable. Because we predicted differential effects over time between groups, we conducted planned contrasts from these models assessing the effect of time within condition, and differences between conditions at each time point (test-wise alpha = 0.05)

Due to the number of behavioral variables, and substantial correlations between them, we conducted principal components analyses (PCA) to derive a smaller set of behavioral measures for statistical modeling. Because the postural and locomotion variables were coded for subjects in both the control and HAI conditions, we performed a PCA with these measures including data from all subjects. For subjects in the HAI condition, we conducted a second PCA including measures related to social interaction. The postural variable “lying (supine)” was observed predominantly in conjunction with stomach petting in the HAI condition and was rarely observed in the control group (median duration in control group = 0). Thus, we omitted this measure from the PCA with variables related to locomotion, and included it in the PCA with variables related to social interaction. To limit the impact of outliers and skew, all behavioral variables were Yeo-Johnson transformed (Yeo & Johnson, 2000), centered, and scaled prior to fitting the PCA. To determine the number of components to retain, we conducted parallel analysis (Horn, 1965) comparing eigenvalues from the actual data to randomly resampled (with replacement, and dimensions equal to those of the original data) and simulated data (random data from a normal distribution). These analyses suggested retention of two components for both the locomotion and social interaction PCAs. To assess relationships between these behavioral variables and changes in OT/AVP across the experiment, we fitted linear models predicting the log transformed percent change in OT/AVP concentration as a function of fixed effects for sex (male, female) and scores from the first two principal components. To accommodate negative values in the dependent measure, we first added a constant to all values (the absolute value of the smallest observation + 1) prior to log transformation. Because the control group did not have component scores for the social interaction variables, models for this group included only sex and principal component scores for the PCA on locomotion/postural variables. Coefficients for fixed effects were tested with likelihood ratio tests comparing the full model, to nested models with the removal of individual terms. All statistical models used a test-wise alpha threshold of 0.05.

## Results

### Between group OT/AVP effects

Figure 1 shows the mean OT/AVP concentrations across time in the HAI and control groups. The results of planned contrasts comparing OT/AVP concentrations between groups, and assessing within group changes over time, are shown in Table 3.

**Figure 1.**
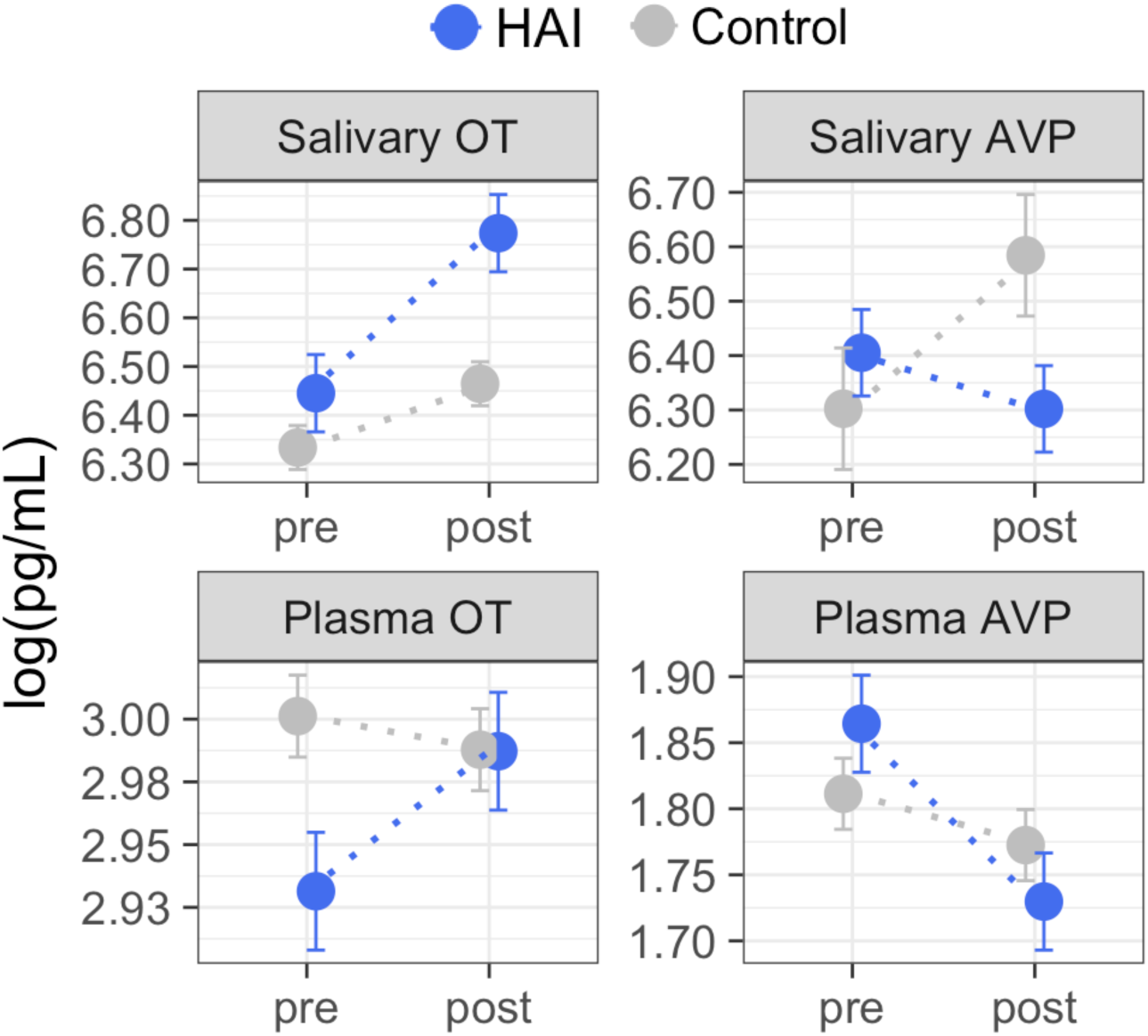
Group means for oxytocin (OT) and arginine vasopressin (AVP) measured in plasma and saliva before and after Experiment 1. Using between-subjects design, dogs were assigned to conditions consisting of 10 minutes of affiliative interaction with a human (HAI, N = 19) or 10 minutes of rest without social contact with the experimenter (Control, N = 19). Dogs in the HAI condition exhibited significant increases in both salivary and plasma OT, and a significant decrease in plasma AVP. Dogs in the control group exhibited no significant changes in salivary or plasma OT, but showed increases in salivary AVP across time. Error bars represent the within-subjects standard error (Cousineau, 2005) and should be interpreted with regard to within-group, but not between group differences. The results of planned between-group, and within-group contrasts are reported in Table 3.

**Table 3.** Results of planned contrasts between the HAI and control groups at each time point, and across time within groups. All tests had 1 degree of freedom. For group comparisons, β indicates the mean difference (HAI – Control). For change over time, β reflects the mean change from pre to post.

|  | Group Comparison (HAI vs. Control) |  |  |  |  |  | Within Group Change (Pre vs. Post) |  |  |  |  |  |
| --- | --- | --- | --- | --- | --- | --- | --- | --- | --- | --- | --- | --- |
|  | Pre-test |  |  | Post-test |  |  | HAI group |  |  | Control Group |  |  |
| | $\beta$ | SE | p | $\beta$ | SE | p | $\beta$ | SE | p | $\beta$ | SE | p |
| Salivary OT | 0.11 | 0.13 | 0.39 | 0.31 | 0.13 | 0.02 | 0.33 | 0.09 | < 0.01 | 0.13 | 0.09 | 0.16 |
| Salivary AVP | 0.12 | 0.21 | 0.58 | -0.27 | 0.21 | 0.20 | -0.10 | 0.13 | 0.44 | 0.28 | 0.13 | 0.04 |
| Plasma OT | -0.07 | 0.07 | 0.35 | 0.00 | 0.07 | 0.97 | 0.06 | 0.03 | 0.05 | -0.01 | 0.03 | 0.64 |
| Plasma AVP | 0.05 | 0.07 | 0.49 | -0.05 | 0.07 | 0.47 | -0.13 | 0.05 | < 0.01 | -0.04 | 0.05 | 0.39 |

For salivary OT, planned comparisons revealed that whereas the control and experimental groups had comparable salivary OT levels at baseline, dogs in the experimental group had significantly higher post-test salivary OT than controls. Similarly, salivary OT exhibited a significant increase from baseline to post-test in the HAI, but not the control group (Figure 1; Table 3). For salivary AVP, there were no between group differences at either time point, however the control group exhibited a significant increase in salivary AVP whereas the HAI group did not (Figure 1; Table 3). Plasma OT did not differ between groups at either time point, however, only the HAI group exhibited a modest, but significant, increase in plasma OT over the course of the study (Figure 1; Table 3). TLastly, there were no group differences in plasma AVP at either time point, however the HAI group exhibited a significant decrease in plasma AVP across time, whereas the control group did not (Figure 1; Table 3).

### Behavioral predictors of changes in OT and AVP

We retained two components from the PCA including variables related to locomotion and posture (LP), which collectively explained 80% of variance in these behaviors. Variable loadings from this model are shown in Table 4. The first component (LP-PC1, 50% variance explained) had strong positive loadings for locomotion and upright posture and moderate negative loadings for lying in in a prone position. The second component (LP-PC2, 30% variance explained) had a strong positive loading for sitting and moderate negative loadings for locomotion, upright posture, and lying (prone).

**Table 4.** Variable loadings from principal component analyses including variables related to locomotion and posture (all subjects) and social interaction (HAI condition only).

|  |  | Loadings |  |
| --- | --- | --- | --- |
|  |  | PC1 | PC2 |
| Locomotion / Posture<br>PCA (all subjects) | locomotion | 0.62 | -0.17 |
|  | upright | 0.64 | -0.16 |
|  | sitting | -0.02 | 0.83 |
|  | lying (prone) | -0.44 | -0.50 |
| Social Interaction PCA<br>(HAI subjects) | physical contact | -0.62 | 0.23 |
|  | play | 0.55 | 0.32 |
|  | licking | 0.32 | 0.71 |
|  | lying (supine) | -0.46 | 0.58 |

For dogs in the HAI condition, we retained two components from the PCA with variables relating to social interaction (SI), which collectively explained 81% of variance in these behaviors. Variable loadings from this model are shown in Table 4. The first component (SI-PC1, 53% variance explained) was loaded positively by play and licking, and negatively by physical contact and lying (supine). The second component (SI-PC2, 28% variance explained) was loaded positively by all four social interaction variables (physical contact, play, licking, and lying supine).

Associations between changes in OT/AVP concentrations and behavior during the test are shown in Table 5. In the HAI condition, the percent increase in salivary OT was positively associated with SI-PC2 scores, which reflect longer durations of physical contact, play, licking, and lying (supine) with the stomach exposed to the experimenter. Despite this positive association with salivary OT, there were no associations between changes in plasma OT concentrations and any of the behavioral variables (Table 5). There were also no associations between changes in salivary AVP and any of the behavioral variables, however, SI-PC2 scores were both negatively related to the percent change in plasma AVP. On average, subjects in the HAI group exhibited a 10% decrease in plasma AVP across the study. However, subjects with SI-PC2 scores in the upper 50^th^ percentile (high levels of affiliative behavior with experimenter) exhibited a larger decrease in plasma AVP (mean change = -18%, SEM = 6.89%) than subjects with SI-PC2 scores in the lower 50^th^ percentile (mean change = -4%, SEM = 4.87). Within the HAI group, LP-PC2 scores were also negatively associated with the change in plasma AVP. Thus, subjects who engaged in more sitting, and less standing, active locomotion and lying (prone) exhibited larger decreases in plasma AVP across time. Within the control group, there were no significant associations between any of the behavioral measures and changes in peptide concentrations, in saliva or plasma (Table 5).

**Table 5.** Associations between changes in hormone concentrations in plasma and saliva and behavior during the test. PC = principal component. Refer to Table 4 for behavioral variables loading each component. β represents the association between a one unit increase in PC scores, and the log transformed percent change in hormone concentrations across time.

| matrix / hormone |  | Social Interaction - PC1 |  |  | Social Interaction - PC2 |  |  | Locomotion & Posture - PC1 |  |  | Locomotion & Posture - PC2 |  |  |
| --- | --- | --- | --- | --- | --- | --- | --- | --- | --- | --- | --- | --- | --- |
|  |  | β | χ² | p | β | χ² | p | β | χ² | p | β | χ² | p |
| HAI group | Salivary OT | -0.60 | 1.55 | 0.21 | 0.61 | 4.37 | 0.04 | 0.72 | 2.70 | 0.10 | 0.22 | 0.60 | 0.44 |
|  | Salivary AVP | -0.02 | 0.00 | 0.97 | 0.16 | 0.22 | 0.64 | 0.10 | 0.04 | 0.84 | 0.18 | 0.26 | 0.61 |
|  | Plasma OT | -0.26 | 0.44 | 0.51 | 0.04 | 0.04 | 0.84 | 0.25 | 0.51 | 0.48 | 0.18 | 0.60 | 0.44 |
|  | Plasma AVP | -0.15 | 0.22 | 0.64 | -0.49 | 6.15 | 0.01 | 0.21 | 0.52 | 0.47 | -0.40 | 3.96 | 0.05 |
| Control group | Salivary OT | - | - | - | - | - | - | -0.06 | 0.07 | 0.80 | -0.05 | 0.03 | 0.85 |
|  | Salivary AVP | - | - | - | - | - | - | -0.36 | 1.80 | 0.18 | 0.20 | 0.35 | 0.56 |
|  | Plasma OT | - | - | - | - | - | - | 0.12 | 0.89 | 0.35 | 0.13 | 0.58 | 0.45 |
|  | Plasma AVP | - | - | - | - | - | - | 0.13 | 2.49 | 0.12 | -0.04 | 0.15 | 0.70 |

## Discussion

We assessed changes in dog salivary and plasma OT and AVP in response to HAI, or a control condition. Dogs in the HAI group exhibited increases in both salivary and plasma OT, whereas neither of these effects were observed in the control group. Dogs in the HAI group exhibited a decrease in plasma (but not salivary) AVP across time, whereas dogs in the control group exhibited an increase in salivary AVP, with no significant changes in plasma AVP. Within the HAI group, the extent of the increase in salivary OT, and the decrease in plasma AVP were predicted by the degree of affiliative contact between the dog and experimenter. Therefore, both the between and within group differences are most likely attributable to the social interactions between the experimenter and dog, and are unlikely to be accounted for by alternative explanations such as a stress response following the initial sample collection.

Although we observed a significant increase in both plasma and salivary OT, the effect was much more pronounced in saliva, echoing the findings of our pilot study. There are at least three reasonable explanations for this finding. First, plasma OT responses can occur extremely rapidly, and may be most evident within 90 seconds of a triggering event (Jonas et al., 2009). In contrast, time series analyses suggest a delay in the transfer of hormones from plasma to saliva, and hormonal peaks in saliva have been documented to occur ~10 minutes after those in blood (Hernandez et al., 2014). Although the timing, and specific mechanisms through which OT and AVP reach saliva in dogs are unknown, it is probable that changes in salivary concentrations lag behind those in plasma, despite the fact that changes in salivary OT occur quickly relative to other salivary hormones (de Jong et al., 2015). Because this study was designed predominantly for salivary measures, we collected samples at the time point characterized by the highest salivary OT levels in our pilot study. Therefore, is it possible that the largest changes in plasma OT occurred quickly, as has been documented in previous studies (Handlin et al., 2011). Nonetheless, we detected a small but significant increase in plasma OT within the HAI group, suggesting similar effects in both blood and saliva.

Second, whereas OT can be measured without interference in non-extracted dog saliva, other components of dog plasma interfere with OT ELISAs, necessitating additional preparatory procedures such as solid phase extraction for the measurement of free OT (MacLean, Gesquiere, Gruen, et al., Submitted). Although extraction procedures can eliminate interfering substances (e.g. albumin proteins), they are often characterized by poor recovery of the target analyte, and thus may eliminate ‘signal’ as well ‘noise’. Unpublished data from our laboratory suggest that common extraction procedures fail to recover substantial amounts of free OT in plasma, despite performing well with kit standards. Therefore, in addition to benefits related to welfare, by virtue of not requiring extraction procedures, salivary measures of OT also have methodological advantages relative to blood sampling.

Third, it has recently been demonstrated that OT rapidly binds to other molecules in plasma, and bound OT is likely to evade detection (Brandtzaeg et al., 2016). Although the binding properties of OT in saliva are unknown, it is possible that OT remains free – and consequently detectable – in saliva, more so than in plasma. This possibility is supported by the high concentrations of OT that we observed in saliva relative to plasma, an effect that persists even with solid phase extraction of dog saliva samples (MacLean, Gesquiere, Gee, et al., Submitted).

In addition to these effects on OT, dogs in the HAI condition also exhibited a decrease in plasma AVP, whereas dogs in the control group exhibited an increase in salivary AVP. Because AVP can activate the hypothalamic-pituitary-adrenal (HPA) axis (Aguilera & Rabadan-Diehl, 2000; Scott & Dinan, 1998), one plausible explanation is that these differences reflect a short-term reduction in stress reactivity in the HAI group, and increased stress reactivity in the control group. This possibility is consistent with many other studies documenting that positive social interactions can buffer stress responses (DeVries, Glasper, & Detillion, 2003) – including interspecies interactions (Schöberl et al., 2016) – whereas social isolation can have the opposite effect. Although dogs in the control group remained in the same room with the experimenter, they were physically separated from him, and this separation may have imposed a mild psychological stressor. Alternatively, it is possible that the initial blood and saliva collection imposed an acute stressor in both groups, but that this event was socially buffered by OT in the HAI condition. The latter possibility is consistent with many studies documenting OT’s ability to supress HPA activity (Carter & Altemus, 1997). Although we cannot distinguish between these, and other explanations at present, future studies will benefit by exploring the dynamics between OT, AVP and HPA activity in the context of HAI. Additionally, the differential changes in AVP between groups were observed in different matrices (increased salivary AVP in the control group, decreased plasma AVP in the HAI group), and the explanation for this phenomenon remains unknown.

Lastly, in addition to the between group differences that we observed, within the HAI group the extent of the increase in salivary OT, and decrease in plasma AVP, were predicted the nature of interactions between the dog and experimenter. Specifically, dogs who engaged in higher levels of physical contact, play, licking and lying in a supine position with the experimenter exhibited the largest increases in OT, and decreases in AVP. Thus, as in previous studies, changes in OT and AVP concentrations depended not only on human contact, but also on the nature of these interactions (Nagasawa et al., 2015; Rehn et al., 2014).

Collectively, our results corroborate previous findings suggesting that OT responds dynamically to HAI, and provide the first data on HAI’s effects on AVP in dogs. Notably, although HAI-related effects on OT have also been observed in humans, we are not aware of any studies examining AVP in humans in the context of interaction with animals. Given that we observed significant decreases in dogs’ plasma AVP concentrations following HAI, we expect that future studies will benefit by incorporating this measure in humans as well. Collectively, these studies suggest that salivary measures of OT and AVP provide noninvasive biomarkers which respond to aspects of affiliative social behavior, and provide researchers with a new set of tools for exploring the roles of OT and AVP in the biology of human-animal interaction.

## Acknowledgements

We are grateful to Brenda Kennedy and staff at Canine Companions for Independence for their help with this research. We thank Hossein Nazarloo, Martina Heer, and Taichi Inui for helpful conversations, and Susan Alberts for allowing us to work her laboratory. We gratefully acknowledge support from the WALTHAM^®^ Centre for Pet Nutrition, which funded this research, and the Stanton Foundation, for a Next Generation Canine Research Fellowship to ELM.

